# A graphical, interactive and GPU-enabled workflow to process long-read sequencing data

**DOI:** 10.1101/2021.05.11.443665

**Authors:** Shishir Reddy, Ling-Hong Hung, Olga Sala-Torra, Jerald Radich, Cecilia CS Yeung, Ka Yee Yeung

## Abstract

We present a graphical cloud-enabled workflow for fast, interactive analysis of nanopore sequencing data using GPUs. Users customize parameters, monitor execution and visualize results through an accessible graphical interface. To facilitate reproducible deployment, we use Docker containers and provide an Amazon Machine Image (AMI) with all software and drivers pre-installed for GPU computing on the cloud. We observe a 34x speedup and a 109x reduction in costs for the rate-limiting basecalling step in the analysis of blood cancer cell line data. The graphical interface and greatly simplified deployment facilitate the adoption of GPUs for rapid, cost-effective analysis of long-read sequencing.

## MAIN TEXT

Cancer is a leading cause of mortality worldwide with the highest burden of death affecting lower- and middle-income countries [1]. Delays in medical care from the inability to detect cancer earlier are a key component contributing to higher morbidity, poor response to treatments and lower survival [2]. Advances in molecular diagnosis have enabled detection of specific driver mutations that can be essential for prognosis, monitoring, and targeted therapy [3-5]. Examples of such “precision medicine” include the BCR-ABL fusion gene in chronic myeloid leukemia (CML), PML-RARA fusions in acute promyelocytic leukemia (APL) and FLT3 mutations in acute myeloid leukemia (AML) [6-9]. Potentially targetable mutations are also found in solid tumors, such as renal cell carcinoma [10-12]. Currently, detection of fusion genes by chromosomal analysis requires highly specialized laboratories. Chromosomal analyses may not have the precision necessary to identify the specific breakpoint in a patient or provide the sequence of the flanking segments around the fusion gene to allow for downstream development of patient specific monitoring assays. Routine PCR based assays can be rapid, but a priori knowledge of the fusion breakpoints is required and when fusions involve large intronic regions RNA input are generally needed. Thus, turn-around times for these methods can take three days to two weeks [13, 14]. To capitalize on the potential of precision medicine, faster analysis of sequencing data is needed to improve the potential of molecular-assisted cancer diagnoses [15].

For cancer management, next generation sequencing (NGS) has many limitations, such as phasing errors, mis-mapping from short reads, strand bias and amplification errors causing irregular variant allele frequency, and bioinformatically-challenging repetitive sequences [16, 17]. In contrast, long-read sequencing technology, such as Oxford Nanopore Technologies (ONT), generates continuous sequences up to a few megabases in length at this time [18]. Nanopore sequencing provides both sequencing and phasing information because the read lengths are generally very long in comparison to NGS (NGS= 150 to 250 bp, nanopore= generally >2kb to 200,000kb or longer), therefore it will not incur the same artifacts and mis-mapping errors [19]. Unlike NGS, which takes an average of three days to complete sample processing and library preparation plus one additional day for sequencing, nanopore sequencing can directly sequence DNA at amazingly fast speeds [20, 21]. Thus, long-read sequencing technologies hold promise in overcoming the current diagnostic gap in cancer research. Computational methods and software tools tailored for long-read sequencing data are essential to enable use of this emerging and promising technology [22, 23]. In nanopore sequencing, electrical current alterations are recorded as different bases traverse the pore opening. Basecalling, which translates the signal (stored as fast5 files) into a sequence of base pairs is the key step determining accuracy of the sequencing experiment. The initial conversion is followed by error correction and data polishing to obtain the final sequence [23]. Basecalling is computationally expensive and a rate-limiting step in the analysis of nanopore data. For NGS data, aligning reads is the rate limiting step due to the greater number of reads and the absence of a separate basecalling step. Deep learning neural network models have been applied to basecalling to increase the accuracy [24]. With standard CPU processing, these methods are prohibitively slow and require large numbers of computational cores operating in parallel to be practical. Graphics processing units (GPUs) can be used to accelerate the analysis but require specialized hardware and software. Hardware in the form of GPU instances are available on public cloud services such as Amazon Web Services (AWS). However, virtual machine instances do not come with the drivers or Compute Unified Device Architecture (CUDA) libraries installed. In addition, the versions of these drivers and libraries must be carefully matched to the software being executed.

Nanopore sequence is fast and cost effective in terms of data collection, with a $90 USB Flongle attached to a laptop acquiring data in a couple of hours. Sequencing is potentially accessible to a broad range of biomedical scientists who do not have access to a traditional sequencer. A challenge in democratizing long-read sequencing technology is the difficulty of the processing of the data which requires command line tools and technically difficult installation of software, libraries and drivers. There is a lack of *graphical* bioinformatics software tools which can efficiently process the raw nanopore reads, and *interactive* visualizations for interpretations of results. An example of a command-line workflow is the MasterOfPores pipeline that performs pre-processing and analysis (prediction of RNA modifications and estimation of polyA tail lengths) of long-read data [25]. MasterOfPores is a workflow using the NextFlow framework [26], a script-based engine that requires programming experience to deploy and modify. While MasterOfPores uses software containers for most of its components, it does not provide a container for the key basecalling step which requires the most setup and configuration to operate with GPU computing. In addition, MasterOfPores does not include the product-grade basecaller Guppy [27], which is available to ONT customers via their community site [28] and cannot be distributed in a container.

We present a graphical cloud-enabled containerized workflow for fast, interactive analysis of nanopore data using GPUs. Specifically, we extended the Biodepot-workflow-builder (Bwb) [29] to provide a modular and easy-to-use graphical interface that allows users to create, customize, execute, and monitor bioinformatics workflows. **Figure 1** shows screenshots of the platform. The workflow consists of modules to download the data and genome files, basecallers Guppy [27] or Bonito [30], minimap2 [31] for sequence alignment, and the Integrated Genome Viewer (IGV) [32, 33] for visualization of the BAM files. We provide containers for both the ONT proprietary Guppy [27] and the ONT open-source Bonito [30] basecallers. In accordance with the licensing constraints of Guppy, we provide a containerized setup module that creates the Guppy container locally when the user provides the download URL from the ONT community site. To facilitate deployment, we use Docker containers for all modules (including the Bwb platform) and provide an Amazon Machine Image (AMI) with all software and drivers pre-installed for GPU computing on the cloud.

**Figure 1.**
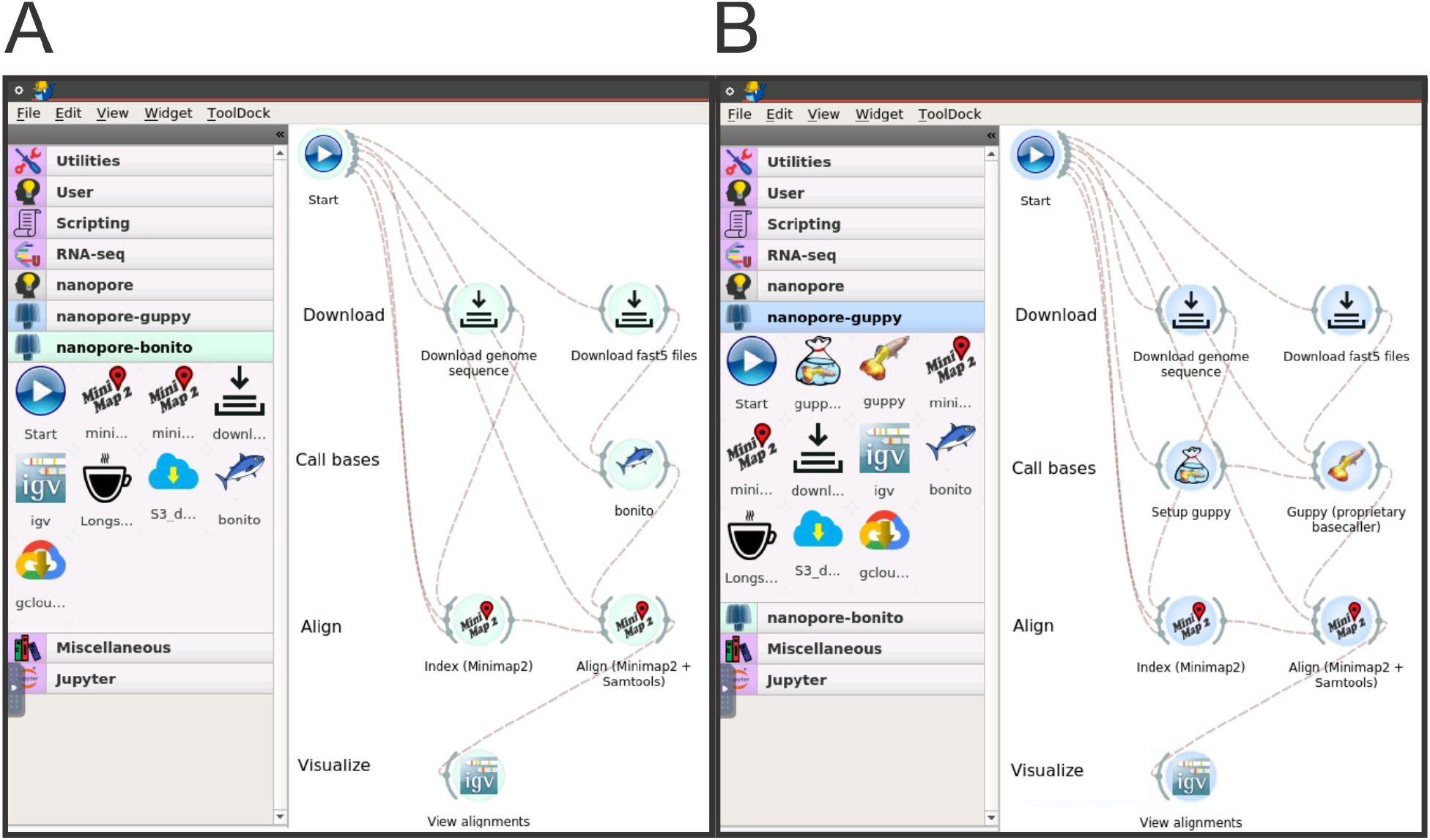
Screenshots of our interactive GPU workflow which uses the Biodepot-workflow-builder platform. Panel A is a screenshot of the workflow using the open-source Bonito basecaller. Panel B is a screenshot of the workflow using the proprietary Guppy basecaller. Both basecallers use GPUs. For the Guppy workflow, the user enters the URL for the Oxford Nanopore Technology Guppy installation package which is then used to create a container to execute Guppy. The other steps in the two workflows are identical, consisting of data download, alignment and visualization. Each of these steps are performed by software modules encapsulated in Docker containers and represented by the graphical widgets. Lines connecting the widgets indicate flow of data between the modules. The user double clicks on the Start widget, enters the necessary parameters into the forms and presses a graphical start button to start the workflow. Double-clicking on a widget brings up a point-and-click interface for users to enter parameters, monitor results and control execution of the associated workflow module. Unlike other workflow execution platforms, the Biodepot-workflow-builder supports modules with interactive graphics. This is leveraged in this workflow to automatically open the final BAM files in the Interactive Graphics Viewer (IGV) which we use to check for diagnostic translocation breakpoints in our cell-line data. The execution time of the basecallers Guppy and Bonito on GPU-enabled machines using the NB4 cell line averaged 75.9 seconds (standard error 0.4) and 948.2 seconds (standard error 1.7) on an AWS g4dn.4xlarge GPU instance. For comparison, the CPU version of Guppy averaged 2551.8 seconds (standard error 22.4) on a AWS virtual machine instance (C5d.18xlarge) using 72 vCPUs.

We demonstrate the benefits of our tools by comparing the execution time of the basecalling step with and without GPU computing. The runtime of Guppy is reduced from over 42 minutes to just over 1 minute using GPU computing, representing a 34x speedup and a 109x reduction in cloud computing costs (see **Methods** for details).

Applications of nanopore sequencing coupled with our workflows in cancer diagnostics are shown in **Figure 2**. We can reliably detect fusion genes from DNA sequences using cell lines (NB4, K562, KCL22, and KU812) with known fusion genes. This provides an advancement to molecular diagnostics by its ability to detect specific breakpoints even if there is a large intronic region between the fusion genes of interest. DNA sequencing of NB4 on a comparatively low cost ($90) Flongle nanopore device, (Oxford, UK), confirmed fusion gene sequencing spanning the *PML* and *RARA* genes (**Figure 2a**). DNA sequencing of K562 on a Flongle was able to capture the fusion gene spanning the *BCR* and *ABL1* genes, as well as capturing the intervening large *ABL1* intronic region 1 where the specific breakpoint occurs (**Figure 2b**).

**Figure 2.**
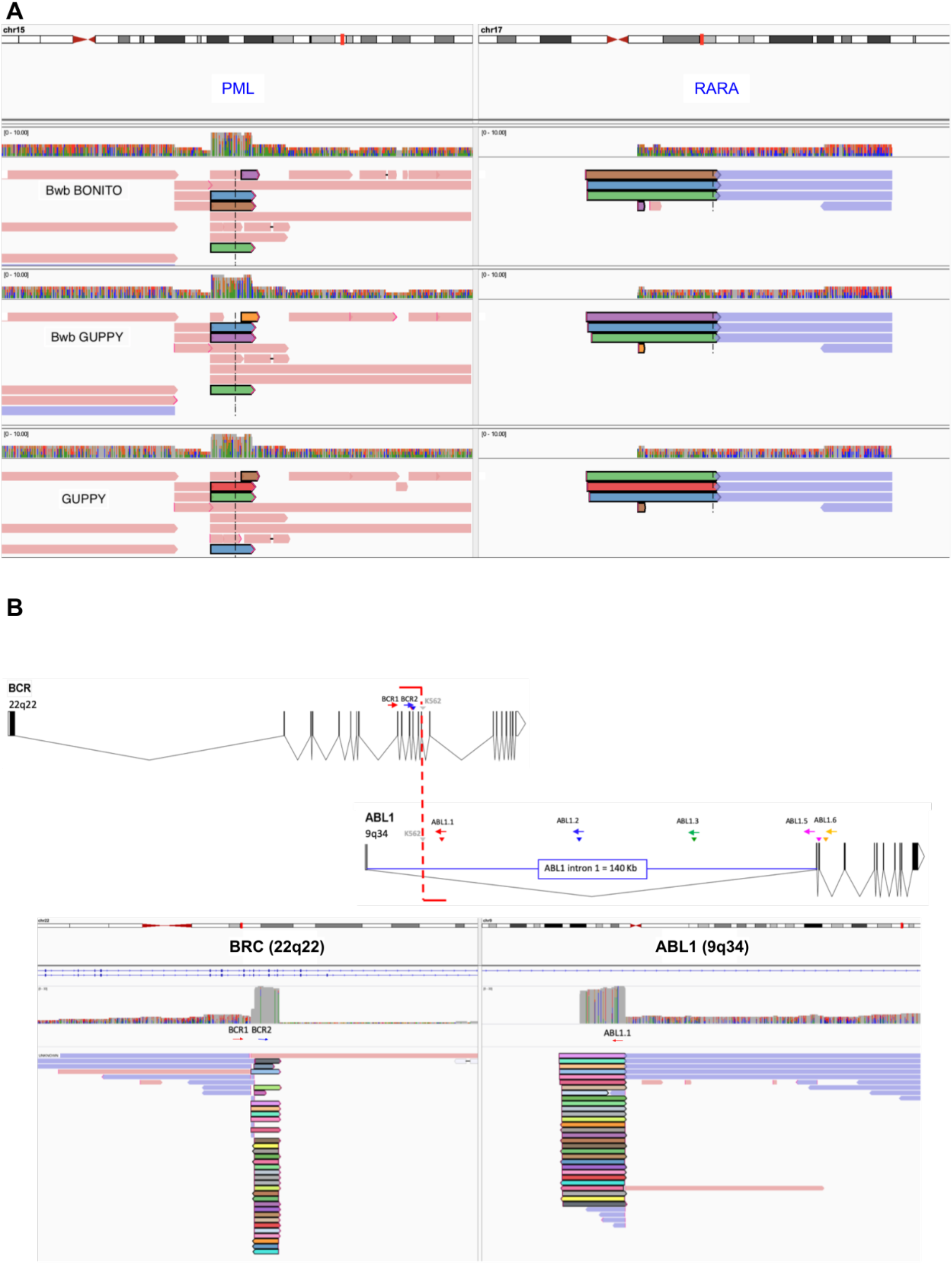
A. IGV viewer alignment on the PML and RARA genes of a Flongle Nanopore generated sequences for the NB4 cell line. Library was generated from DNA with a PCR-free enrichment protocol using CRISPR guides targeting PML and RARA genes. The top panel shows the alignment for the reads processed with the Bonito basecaller and minimap2 aligner in the Bwb. The middle panel shows the alignment of reads processed with the Guppy basecaller and minimap2 aligner in the Bwb workflow. The bottom panel shows reads processed in a manual step-by-step workflow using the Guppy flipflop basecaller and minimap2 aligner. Reads with PML-RARA breakpoint are colored to highlight the fragment aligned to PML and RARA. **Figure 2B**. Genomic BCR-ABL1 breakpoint identified in the K562 cell line by long-read sequencing. Schematic representation (generated with http://wormweb.org/exonintron) shows the breakpoint captured with our amplification-free enrichment protocol and long-read sequencing. The breakpoint is represented in the upper graphic by the red vertical line, and the location of the sequence specific guides is marked by colored arrows. ABL1 intron 1 spans 140Kbs. In the lower panel, nanopore sequence alignments in IGV show sequences partially aligned to BCR and ABL1. Reads with BCR-ABL1 breakpoint are colored to highlight the same read is partially aligned to BCR and ABL1.

A potential advantage of long-read sequencing is lower costs in comparison to standard NGS. This is dependent on manufactures of nanopore products to maintain use of comparatively inexpensive parts and designs for smaller portable devices. We used low-cost Flongles to detect the PML-RARA and BCR-ABL1 fusions. In this work, we demonstrate the ability to detect fusions on nanopores devices that read DNA at speeds faster than 1 nt/μs. We present not only interactive software tools that make analyses of Nanopore data accessible to biomedical and clinical scientists, but also demonstrate these analyses are efficient and cost effective using GPU computing. Most importantly, we have illustrated the applicability of our workflow using cell line data in a rapid, cost effective assay to detect fusions through our integrated bioinformatics workflow which can process raw nanopore reads, perform basecalling, support graphical output and visualizations for interpretations of results on an easy user interface. This allows the capability to identify and confirm pathognomonic fusion genes such as BCR-ABL1 in CML or PML-RARA in APL. This methodology can potentially both define specific breakpoints important in treating and tracking a patients’ specific disease, but also allow multiplex phasing to identify and follow multiple mutations in the same patient. Future directions include optimizing the assay for faster library preparation, sequencing times, and analysis to enable fusion detection to less than a day. Vast improvements to turnaround time in the laboratory and an accessible, efficient informatics workflow that enables most molecular technologists and pathologists to implement long-read sequencing into current clinical pathology workflows will advance the field of molecular pathology beyond NGS. These small portable, low-cost devices, together with integrated bioinformatics support, will allow for rapid diagnostics assisting point-of-care clinical decision making.

## METHODS

### Data generation using cell lines

5,000 ng of DNA from two cell lines, NB4 and K562, was dephosporylated and cut with Cas9 enzyme complexed with RNA guides designed to target the genes involved in their respective translocations (BCR and ABL1 for K562, and PML and RARA for NB4). Nanopore adapters were then ligated to the newly created DNA ends and the library was loaded onto a flow cell or Flongle and sequenced using a MinIOn Mk1B.

#### Integrated and interactive workflow in the Biodepot-workflow-builder (Bwb)

Biodepot-workflow-builder (Bwb) [29] provides a modular and easy-to-use graphical interface for reproducible execution, customization and interactive visualization of the nanopore pipeline. Graphical widgets representing Docker containers that execute modular tasks are graphically linked to define bioinformatics workflows that can then be reproducibly deployed across different local and cloud platforms. New widgets (modules) can be added without writing code. In this work, we created new widgets in the Bwb for basecalling using Guppy [27] and Bonito [30], alignment using minimap2 [31], visualization of the resulting BAM files using the Integrated Genome Viewer (IGV) [32, 33]. Each of these widgets call a Docker container in the backend. Users can adjust input parameters of each widget using an intuitive form-based user interface, and check intermediate results using a console. A key characteristic of Bwb is integrated support for graphical output, enabling *interactive* tools such as Jupyter notebooks, spreadsheets, and visualization tools to be included in the workflow. We leverage this graphical output support feature of the Bwb to create an integrated, interactive workflow by integrating the IGV to visualize resulting BAM files. Our workflows can be shared using a Bwb native format or exported as shell scripts. In addition, the Bwb is distributed as a Docker container that can be easily deployed on *any local or cloud platform*.

### GPU software versioning and compatibility ensured by using containers and providing an AMI with drivers and software

GPU-enabled executables require additional software layers to use the GPU hardware. For NVIDIA hardware, CUDA software provided by the manufacturer is necessary to perform general computations on the GPU. AMD GPUs use a different and incompatible set of software for their hardware. In addition, low level libraries (drivers) are required to communicate with the GPU card. Additional language and operating system dependent libraries and headers may also be required to integrate the CUDA software. All these layers of software interact with each other and as a result, compatibility is version dependent and even sensitive to the method of installation. Components installed using scripts may be incompatible with components installed using package managers.

An example of the complexities involved in deploying GPU software is the open source Bonito basecaller. The current version of the Bonito caller will not install if one follows the instructions on the Github [30]. This is due to dependency incompatibilities in the current Python PyTorch [34] packages and CuPy [35] libraries. We were able to install Bonito v0.38 by downgrading from CUDA 11.3 to CUDA 10.2, cuDNN 7, CuPy 10.2. and downgrading PyTorch from 1.8.0 to 1.7.1. By providing a Docker container we can ensure that Bonito is deployed in this compatible environment without the need for the user to install the exact versions of the software. However, this is not sufficient, the user still needs to install the compatible drivers and CUDA software on their host machine. This is true even for GPU instances on Amazon cloud. AWS does provide the basic Linux operating systems but requires that users install their own drivers and libraries. There are commercial distributions on the AWS marketplace that provide support but we could find nothing among the free community offerings. As a resource for the nanopore research community, we therefore have provided a public freely available AMI for use with AWS GPU virtual machine instances with versions of drivers and libraries that we have tested with our workflows. The combination of AMI and containerized modules eliminates the arcane installation steps and makes the nanopore GPU software accessible to a broad audience in the biomedical and clinical community.

### Benchmarking basecallers

We compared the execution time of the basecallers Guppy [27] and Bonito [30] on GPU-enabled machines using the NB4 cell line data. We also measured the execution time of running Guppy using CPU. We repeated our empirical experiments four times and observed that Guppy GPU achieved the fastest average time at 75.9 seconds (1.3 minutes) with standard error of 0.4. Bonito GPU finished basecalling in 948.2 seconds (15.8 minutes) on average with standard error of 1.7. Guppy CPU achieved the slowest average time at 2551.8 seconds (42.5 minutes) with standard error of 22.4.

### Experimental setup for benchmarking

Guppy GPU was benchmarked on an AWS g4nd.4xlarge virtual machine instance with a NVIDIA Tesla T4 GPU using the dna_r9.4.1_450bps_hac.cfg configuration file and template_r9.4.1_450bps_hac.jsn model file provided by ONT. Bonito GPU was also benchmarked on the same instance using the provided dna_r9.4.1 model file and the default settings (chunk size of 4000 and batch size of 32). Guppy CPU was benchmarked on a c5d18xlarge instance with 72 vCPUs, 72 threads/basecaller, and 1 basecaller. All benchmark experiments were based on 4 runs.

### Estimation of costs of basecalling step

A large selection of virtual machine instance types with different pricing structures are available on AWS [36]. We conducted our empirical experiments using the C5 and G4 EC2 (Elastic Compute Cloud) instances, designed for compute-intensive workloads and GPU computing respectively. In the us-east-2 region, the on-demand pricing of the AWS c5d.18xlarge EC2 instance, with 72 vCPUs and 144GB memory, is $3.888 per hour. The pricing for a g4nd.4xlarge, with single GPU, 16 vCPUs and 64GB memory, is $1.204 per hour [36]. The ratio of the costs of CPU vs GPU is the time ratio 2551.8/75.9 multiplied by the pricing ratio 3.888/1.204 which works out 108.6 fold cheaper when the GPU instance is used for basecalling. These cost estimates are based on single samples.

## CODE AVAILABILITY

All code is publicly available and distributed under a custom academic license at https://github.com/BioDepot/nanopore-gpu. A public AMI (ami-0ecb1effab7fcfaa3) is provided with all necessary drivers and software pre-installed. We have also created a demo video on Youtube https://youtu.be/yPhBKjdi8gY

## Authors’ contributions

Conceptualization: J.R., C.Y. and K.Y.Y. Software development and testing: L.H.H. and S.R. Code documentation: S.R. Cloud resources: L.H.H. Computational benchmarking: L.H.H. and S.R. Data generation: O.S. and C.Y. Data analysis: O.S. and S.R. Figure preparation: L.H.H., O.S. and S.R. Writing - original draft preparation: K.Y.Y. and C.Y. Writing - reviewing and editing: All authors. Funding acquisition: J.R., C.Y. and K.Y.Y. The authors read and approved the final manuscript.

## REFERENCES

1. GLOBOCAN 2018: counting the toll of cancer. Lancet 2018, 392:985.

2. Milner DA Jr.,, Holladay EB: Laboratories as the Core for Health Systems Building. Clin Lab Med 2018, 38:1–9.

3. Mehta S, Shelling A, Muthukaruppan A, Lasham A, Blenkiron C, Laking G, Print C: Predictive and prognostic molecular markers for cancer medicine. Ther Adv Med Oncol 2010, 2:125–148.

4. Yeung CCS, Radich J: Predicting Chemotherapy Resistance in AML. Curr Hematol Malig Rep 2017, 12:530–536.

5. Yaghmaie M, Yeung CC: Molecular Mechanisms of Resistance to Tyrosine Kinase Inhibitors. Curr Hematol Malig Rep 2019, 14:395–404.

6. Radich J, Yeung C, Wu D: New approaches to molecular monitoring in CML (and other diseases). Blood 2019, 134:1578–1584.

7. Dohner H, Estey EH, Amadori S, Appelbaum FR, Buchner T, Burnett AK, Dombret H, Fenaux P, Grimwade D, Larson RA, et al: Diagnosis and management of acute myeloid leukemia in adults: recommendations from an international expert panel, on behalf of the European LeukemiaNet. Blood 2010, 115:453–474.

8. O’Donnell MR, Tallman MS, Abboud CN, Altman JK, Appelbaum FR, Arber DA, Attar E, Borate U, Coutre SE, Damon LE, et al: Acute myeloid leukemia, version 2.2013. J Natl Compr Canc Netw 2013, 11:1047–1055.

9. Arber DA, Orazi A, Hasserjian R, Thiele J, Borowitz MJ, Le Beau MM, Bloomfield CD, Cazzola M, Vardiman JW: The 2016 revision to the World Health Organization classification of myeloid neoplasms and acute leukemia. Blood 2016, 127:2391–2405.

10. Xiao X, Garbutt CC, Hornicek F, Guo Z, Duan Z: Advances in chromosomal translocations and fusion genes in sarcomas and potential therapeutic applications. Cancer Treat Rev 2018, 63:61–70.

11. Tretiakova MS, Wang W, Wu Y, Tykodi SS, True L, Liu YJ: Gene fusion analysis in renal cell carcinoma by FusionPlex RNA-sequencing and correlations of molecular findings with clinicopathological features. Genes Chromosomes Cancer 2019.

12. Parker BC, Zhang W: Fusion genes in solid tumors: an emerging target for cancer diagnosis and treatment. Chin J Cancer 2013, 32:594–603.

13. Tsongalis GJ, Al Turkmani MR, Suriawinata M, Babcock MJ, Mitchell K, Ding Y, Scicchitano L, Tira A, Buckingham L, Atkinson S, et al: Comparison of Tissue Molecular Biomarker Testing Turnaround Times and Concordance Between Standard of Care and the Biocartis Idylla Platform in Patients With Colorectal Cancer. American Journal of Clinical Pathology 2020, 154:266–276.

14. Dawson AJ, McGowan-Jordan J, Chernos J, Xu J, Lavoie J, Wang JC, Steinraths M, Shetty S: Canadian College of Medical Geneticists guidelines for the indications, analysis, and reporting of cancer specimens. Curr Oncol 2011, 18:e250–255.

15. VanderLaan PA, Chen Y, DiStasio M, Rangachari D, Costa DB, Heher YK: Molecular Testing Turnaround Time in Non-Small-Cell Lung Cancer: Monitoring a Moving Target. Clin Lung Cancer 2018, 19:e589–e590.

16. Jennings LJ, Arcila ME, Corless C, Kamel-Reid S, Lubin IM, Pfeifer J, Temple-Smolkin RL, Voelkerding KV, Nikiforova MN: Guidelines for Validation of Next-Generation Sequencing-Based Oncology Panels: A Joint Consensus Recommendation of the Association for Molecular Pathology and College of American Pathologists. J Mol Diagn 2017, 19:341–365.

17. Strom SP, Lee H, Das K, Vilain E, Nelson SF, Grody WW, Deignan JL: Assessing the necessity of confirmatory testing for exome-sequencing results in a clinical molecular diagnostic laboratory. Genet Med 2014, 16:510–515.

18. Laver T, Harrison J, O’Neill PA, Moore K, Farbos A, Paszkiewicz K, Studholme DJ: Assessing the performance of the Oxford Nanopore Technologies MinION. Biomol Detect Quantif 2015, 3:1–8.

19. Jain M, Koren S, Miga KH, Quick J, Rand AC, Sasani TA, Tyson JR, Beggs AD, Dilthey AT, Fiddes IT, et al: Nanopore sequencing and assembly of a human genome with ultra-long reads. Nat Biotechnol 2018, 36:338–345.

20. Helmersen K, Aamot HV: DNA extraction of microbial DNA directly from infected tissue: an optimized protocol for use in nanopore sequencing. Sci Rep 2020, 10:2985.

21. Cumbo C, Minervini CF, Orsini P, Anelli L, Zagaria A, Minervini A, Coccaro N, Impera L, Tota G, Parciante E, et al: Nanopore Targeted Sequencing for Rapid Gene Mutations Detection in Acute Myeloid Leukemia. Genes (Basel) 2019, 10.

22. Sedlazeck FJ, Lee H, Darby CA, Schatz MC: Piercing the dark matter: bioinformatics of long-range sequencing and mapping. Nat Rev Genet 2018, 19:329–346.

23. Amarasinghe SL, Su S, Dong X, Zappia L, Ritchie ME, Gouil Q: Opportunities and challenges in long-read sequencing data analysis. Genome Biol 2020, 21:30.

24. Wick RR, Judd LM, Holt KE: Performance of neural network basecalling tools for Oxford Nanopore sequencing. Genome Biol 2019, 20:129.

25. Cozzuto L, Liu H, Pryszcz LP, Pulido TH, Delgado-Tejedor A, Ponomarenko J, Novoa EM: MasterOfPores: A Workflow for the Analysis of Oxford Nanopore Direct RNA Sequencing Datasets. Front Genet 2020, 11:211.

26. Di Tommaso P, Chatzou M, Floden EW, Barja PP, Palumbo E, Notredame C: Nextflow enables reproducible computational workflows. Nat Biotechnol 2017, 35:316–319.

27. Oxford Nanopore Technologies GitHub: Guppy [https://github.com/nanoporetech]

28. The Nanopore Community [https://nanoporetech.com/community]

29. Hung LH, Hu J, Meiss T, Ingersoll A, Lloyd W, Kristiyanto D, Xiong Y, Sobie E, Yeung KY: Building Containerized Workflows Using the BioDepot-Workflow-Builder. Cell Syst 2019, 9:508–514 e503.

30. Bonito: A PyTorch Basecaller for Oxford Nanopore Reads [https://github.com/nanoporetech/bonito]

31. Li H: Minimap2: pairwise alignment for nucleotide sequences. Bioinformatics 2018, 34:3094–3100.

32. Robinson JT, Thorvaldsdottir H, Winckler W, Guttman M, Lander ES, Getz G, Mesirov JP: Integrative genomics viewer. Nat Biotechnol 2011, 29:24–26.

33. Thorvaldsdottir H, Robinson JT, Mesirov JP: Integrative Genomics Viewer (IGV): high-performance genomics data visualization and exploration. Brief Bioinform 2013, 14:178–192.

34. PyTorch [https://pytorch.org/]

35. CuPy: A NumPy-compatible array library accelerated by CUDA [https://cupy.dev/]

36. Amazon EC2 pricing [https://aws.amazon.com/ec2/pricing/]

